# p21^CIP1/WAF1^-mediated partial senescence supports steroidogenesis and therapy resistance in primary prostate cancer cultures

**DOI:** 10.64898/2026.01.15.699666

**Authors:** Emily Toscano-Guerra, Valentina Maggio, José Álvarez-Meythaler, Javier García, Ana Celma, Jacques Planas, Maria Eugenia Semidey, Mercedes Marín, Rosa Somoza, Begoña Mellado, Inés de Torres, Roser Ferrer, Juan Morote, Rosanna Paciucci

## Abstract

**Background:** Progression of prostate cancer to castration-resistant disease is driven by early adaptive mechanisms that remain poorly defined. Increasing evidence suggests that senescence-associated programs and intratumoral steroidogenesis may contribute to therapeutic resistance, but their functional interplay in hormone-naïve disease is unclear.

**Methods:** Hormone-naïve primary prostate cancer cultures (hnPCs) were established from diagnostic biopsies and characterized for proliferation, therapeutic response, senescence markers, and steroidogenic gene expression. Functional relevance of CDKN1A was assessed by knockdown experiments. Public transcriptomic datasets from ETS fusion–negative, AR signaling inhibitor–naïve metastatic castration-resistant prostate cancer (mCRPC) were interrogated for clinical validation.

**Results:** hnPCs exhibited low proliferative capacity yet intrinsic resistance to androgen deprivation therapy and docetaxel. Most cultures displayed coordinated overexpression of CDKN1A/p21^CIP1^ together with steroidogenic enzymes, particularly AKR1C3, alongside p16^INK4a^ expression and SA-β-galactosidase activity, consistent with a partial senescence-like state. CDKN1A depletion selectively reduced AKR1C3 expression and increased proliferation in a subset of cultures, indicating that p21 maintains specific senescence-associated features while permitting adaptive plasticity. Analysis of ETS fusion–negative, ARSI-naïve mCRPC tumors revealed positive correlations between CDKN1A and steroidogenic genes, supporting the translational relevance of this program.

**Conclusions:** These findings identify a previously underrecognized CDKN1A-associated partial senescence program that supports steroidogenic reprogramming, survival, and early therapeutic resistance in aggressive prostate cancer. This adaptive state may predispose tumors to progression toward castration resistance and represents a potential biomarker and therapeutic vulnerability in high-risk, ETS-negative disease.

## INTRODUCTION

Prostate cancer is the most frequent neoplasia and the third leading cause of cancer-related death among men in industrialized countries (1,2). PCa is an androgen-dependent malignancy (3,4), and although radical prostatectomy is effective in eradicating the disease for the majority of patients, those with locally advanced or metastatic cancer face significantly poorer survival (5). Therapies for advanced or metastatic hormone sensitive PCa (mHSPC) include androgen deprivation therapy (ADT), with inhibitors of CYP17A1 (abiraterone) a key enzyme in androgen biosynthesis, or androgen receptor pathway inhibitors (ARPIs) like enzalutamide, apalutamide and darolutamide, which block testosterone production and prevent cancer cells from using remaining testosterone. These approaches are initially effective in significantly reducing tumor burden (6). However, most patients eventually develop resistance and progress to CRPC (7,8).

For patients with CRPC, or with high-risk, high-volume metastatic or non-metastatic disease, treatment options may include combination therapy, radiation, chemotherapy, immunotherapy, and poly-ADP-ribose-polymerase (PARP) inhibitors. Recently, trials testing combinations of enzalutamide and standard ADT have shown a significant reduction in death risk in high-risk recurrent PCa(9). Additionally, a new targeted radiolabeled therapy in patients with metastatic CRPC expressing prostate-specific membrane antigen (PSMA) demonstrated that 177Lu-PSMA-617 (Pluvicto) in combination with ADT plus ARPIs significantly improved progression-free survival (10). Moreover, therapies combining PARP inhibitors with ADT delayed disease progression in patients with aggressive PCa, especially those with *BRCA1/2* mutations(11). Predictive markers of treatment response are needed for early intervention to eliminate aggressive mHSPCs and reduce the risk of resistance. For example, mutations in homologous recombination repair (HRR) genes in patients with mCRPC have enabled clinical implementation of PARP inhibitors as targeted therapies (12).

Uncovering the molecular mechanisms that drive disease progression and therapeutic resistance is a central challenge in PCa research for developing effective and personalized therapies. Various processes have been implicated, including intra-tumoral androgen biosynthesis (13) and cellular senescence(14). To investigate these mechanisms, cancer cell lines and mouse models are commonly used, although the limited number of well-characterized PCa cell lines and the inability of current models to fully replicate tumor phenotypic heterogeneity and native microenvironment continue to present significant challenges (15,16). Traditional cancer cell lines, such as LNCaP, DU145, and PC3, all derived from metastatic lesions, do not adequately reflect the complexity of primary tumor tissue. Although three-dimensional (3D) culture systems, such as spheroids and organoids, more closely mimic tumor architecture, they still lack essential microenvironmental components, limiting their ability to accurately model correlation with patient tumors (17). Patient-derived xenograft (PDX) models preserve tumor architecture and partially reflect clinical responses (18). However, they are constrained by limitations including low engraftment success rates, in several cancers, including PCa (19,20) high cost, and limited representation of early-stage disease(21).

Prostate tumors exhibit significant heterogeneity across clinical, spatial, and morphological dimensions, and their complex genetic landscape poses a considerable challenge to cancer research and personalized therapy (22,23). Primary tumor-derived *ex vivo* cultures have emerged as promising tools that retain the molecular diversity of individual tumors and can predict sensitivity to ADT *in vitro* (24,25). These limitations in existing models, combined with the need to better capture tumor heterogeneity, underscore the potential value of *ex vivo* primary cultures as alternative preclinical platforms that may represent the biology of individual patient tumors.

In this study, we developed *ex vivo* cultures (hnPCs) from patients with aggressive hormone-naïve prostate cancer (HNPC) or metastatic HNPC. To investigate alterations in the steroidogenic pathway associated with tumor aggressiveness, hnPCs were compared to their original diagnostic biopsies, tumor tissues from RP, and established cell lines (LNCaP and DU145). Despite transcriptional shifts that occurred due to *in vitro* conditions, hnPCs preserved key gene expression profiles reflective of their tumors of origin, distinguishing these cultures from conventional cell lines. Notably, a subset of hnPCs exhibited features of partial senescence, characterized by elevated expression of *AKR1C3*, a steroidogenic enzyme involved in intratumoral androgen biosynthesis and *de novo* conversion of adrenal precursors to testosterone. These findings highlight the utility of hnPCs as a valuable alternative model for investigating the mechanisms of prostate tumor progression and therapeutic resistance.

## METHODS

### Patientś selection

The selection of patients for this study was done in collaboration with the Urology Service at the Vall d’Hebron Hospital. Patient characteristics are shown in **Supplementary Table S1**. We collected needle biopsy from prostate tumors of patients with aggressive (untreated) HNPC. Criteria used for patient selection were: Gleason ≥8, PSA >50 ng/mL, and positive DRE. This study was approved by the Medical Research Ethics Committee of Vall d’Hebron Hospital (protocol number: PR(AG) 96/2015).

### Establishment of primary tumor cultures from HNPC tissues

We used five needle biopsies from each patient to establish *in vitro* cultures (hnPCs), while two additional biopsies per patient were selected for nucleic acid extraction. Of the 16 primary tumor cultures we established, 12 were selected for this study. A pathologist confirmed that all primary biopsy cores had a Gleason score of ≥ 8. We initially characterized ten hnPCs (**Supplementary Figure S1**) and later included two additional cultures in our analysis.

### Culture conditions for hnPCs

We established primary cultures using a modified method based on Gao et al. (26). Biopsies were mechanically disaggregated into approximately 1 mm pieces in Basic medium (DMEM-F12, supplemented with amphotericin B 250 ng/µL, gentamicin 10 µg/mL, non- essential aminoacids, penicillin/steptomicin, L-glutamin, and sodium pyruvate). Following centrifugation, tumor pieces were seeded (without enzymatic digestion) onto collagen and poly-D-lysine-coated six-well plates.We used a reduced volume of complete medium (Basic medium with FBS 7%, 10 ng/mL bFGF, 20 ng/mL EGF, 200 ng/mL vitamin A, 200 ng/mL vitamin E, plus 0.6%glucose, 1 mg/mL transferrin, 0.25 mg/mL insulin, 0.097 mg/mL putrescin, 0.3 µM sodium selenite and 100 µM hydrocortisone) for 16 hours to promote cell adhesion. After adding fresh medium, cultures were maintained for 2–3 weeks, replacing the medium every 3 days until cultures reached 70–80% confluence. All cultures were maintained in complete medium at 37°C in a humidified atmosphere with 5% CO2. hnPCs cells were detached using TrypLE (0.5% in PBS) with varies trypsinization times to enrich for epithelial cells and reduce fibroblasts overgrowth (27). Cultures passages were limited to a maximum of 10 to preserve primary cell characteristics and minimize genetic drift.

### Androgen deprivation therapy

Androgen deprivation is the first-line treatment for aggressive PCa. To replicate clinical conditions, androgen-independent (AI) cultures were generated from hnPCs primary cultures by maintaining them in RPMI-1640 medium supplemented with 7% heat-inactivated, charcoal-stripped FBS for several passages. This process mimics the selective pressure of androgen deprivation therapy that patients experience. Surviving cells were detached using TrypLE and replated under the same conditions. We then analyzed AI cultures in comparison to androgen-dependent (AD) cultures maintained in parallel plates under standard conditions.

### Proliferation assay

Cells (2x10^3^ cells/well) seeded on 96-well plates in octuplicates, were grown for 1 to 7 days, then fixed with 4% formaldehyde, washed and stained with 0.5% crystal violet to visualize viable cells. Crystals were dissolved with 15% acetic acid, and optical density was read at 590 nm.

### Cytotoxicity assay

Cells (5x10^3^ cells/well) were seeded in sextuplicates collagen-coated 96-well plates. After 24 hours, different drug concentrations, or vehicle were added and incubated for 72 hours. Then, cells were fixed and processed as described proliferation assays section. For docetaxel and cabazitaxel treatments, aliquots of 1 mg/mL in DMSO solutions were stored at -20°C. These thermolabile drugs were replaced with fresh aliquots daily.

### Western blotting

were performed on PVDF membranes as previously described (28). We used the following primary antibodies: mouse monoclonal anti-p21WAF1 (1:200; Santa Cruz sc-6246), mouse monoclonal anti-MDM2 (1:200; Santa Cruz sc-56154), mouse monoclonal anti-β-Tubulin (1:200; Santa Cruz sc-5274), goat polyclonal anti-actin (1:200; Santa Cruz sc-1616), rabbit polyclonal anti-lamin A/C (1:200; Santa Cruz sc-20681), goat polyclonal anti-Notch1 (1:200; Santa Cruz sc-6014) and rabbit polyclonal anti-AKR1C3 (1:500; Invitrogen PA5-97446). Secondary antibodies were horseradish peroxidase-conjugated polyclonal anti-mouse (1:2,000, Agilent Dako P0260), anti-rabbit (1:2,000, Agilent Dako P0448), or anti-goat (1:2,000, Agilent Dako P0449).

### Real-time quantitative PCR (RT-qPCR)

Total RNA from cells and tissues was extracted using the RNeasy mini kit (QIAGEN). RNA quality was assessed with a 2100 Bioanalyzer (Agilent), and only samples with RIN ≥7 were used for downstream applications. We synthesized cDNA using the NZY M-MuLV first-strand cDNA synthesis kit (NZY Tech) in a 2720 Thermocycler (Applied Biosystems), starting with 500 ng total RNA from biopsies or 1,000 ng from cells/tissues. RT-qPCR was performed on a LightCycler 480 detection system (Roche) using the NZYSpeedy qPCR Probe Master Mix protocol (NZY Tech). Primers from Universal Probe Library (UPL, Roche) or TaqMan probe assay (ThermoFisher Scientific) were used according to availability. Gene-specific primers sequences are listed in **Supplementary Table S2**.

For Sanger DNA sequencing, the cDNA was used for PCR amplification with specific primers for *SPOP, PTEN* and *TP53* genes. The resulting amplicons were sequenced by Sanger (Macrogen Europe Service). For *TMPRSS2-ERG* gene fusion cDNA was amplified in PCR with primers designed to flank both genes, TMPRSS2_E1 FW: 5’-CGCGAGCTAAGCAGGAG-3’, ERG_E6 RV: 5’-CCATATTCTTTCACCGCCCACTCC-3’

### Transient p21-RNA silencing

Predesigned and specific Dicer-Substrate short interference RNAs (DsiRNAs, 27mer siRNA) against *CDKN1A* p21^CIP1^ were obtained from Integrated DNA Technologies, IDT. The kit DsiRNAs (TriFECTa RNAi) contains 3 predesigned DsiRNAs (target-specific), and 3 control DsiRNAs, TYE563 transfection control, HPRT-S1 positive control, and a negative control (scramble). For transfections, LNCaP and hnPCs cells were seeded at ∼50% of confluence in 12-well plates (180 000 cells) or 60 mm plates (800 000 cells), respectively, transfected with *CDKN1A* DsiRNAs (10 nM) or controls, and cultured for 48 hours.

### Immunofluorescence

Cells seeded on sterilized coverslips into 6 well-plates, were cultured for 24 hours, fixed with cold PFA (4%), washed, and permeabilized with Blocking-Permeabilization Buffer (BP) (3% BSA in PBS and 0.25% Triton-X100) for 1h, RT. After washing with BP, coverslips were incubated with p21^CIP1^ antibody (2.0 µg/mL) for 3h then washed, incubated 1h with Alexa Fluor 594 antibody (4 µg/mL) (Invitrogen), then added with phalloidin FITC-conjugated (Sigma-Aldrich) (40 min.). After washing, coverslips were mounted on slides using the ProLong Diamond Antifade Mountant with DAPI (Thermo-Fisher). The primary antibody was omitted in the negative control. Images were acquired using an Olympus FV1000 spectral confocal system integrated with an Olympus IX81 motorized inverted microscope.

### Cell fractionation

Nuclear and cytoplasmic fractions were isolated by differential centrifugation. Cells were seeded in 60 mm plates and grown at 70-80% confluency. All procedures were performed at 4 °C. Cells were washed with PBS, then added with hypotonic buffer (20 mM Tris-HCl pH 7.4, 10 mM KCl, 2 mM MgCl2, 1 mM EGTA, 0.5 mM DTT, 100 µM PMSF, protease and phosphatase inhibitors) and harvested by scraping. After adding Triton-X100 (10%), cells were centrifuged (18,700 × *g*) to separate nuclear (pellet) and cytoplasmic (supernatant) fractions. Supernatants were centrifuged again to eliminate debris. Nuclear fractions were resuspended in isotonic buffer (20 mM Tris-HCl pH 7.4, 150 mM KCl, 2 mM MgCl2, 1 mM EGTA, 0.5 mM DTT, protease and phosphatase inhibitors) and centrifuged (1000 × g). The presence of p21^CIP1^ was analyzed by western blotting with specific antibody.

### Senescence markers

β-Galactosidase expression was analyzed using the CellEvent Senescence Green Detection Kit (Thermo-Fisher) on a Nikon Ts2R FL inverted microscope. Cells were seeded in triplicate 96-well plates (10,000 cells/well) and incubated for 24 hours to allow adherence. Cells were then exposed for 48 hours to different experimental conditions, including basal (untreated), etoposide-treated cells (10 µM, positive control for senescence), transfection controls (dsiRNA TYE563 and scramble siRNA), and siRNAs targeting p21^CIP1^. After washing with 1% BSA in PBS, cells were fixed with 4% paraformaldehyde (PFA), washed again, and incubated with fluoro-X-Gal working solution containing the senescence green probe. Plates were covered with plastic film and incubated in the dark at 37°C for 2 hours without CO₂. Following a PBS wash, images were captured using an Alexa Fluor™ 488/FITC filter set. Cell density was estimated by counting five random non-overlapping fields per well (∼1.23 mm² per field). The expression of *CDKN2A* was also measured as a senescence marker using RT-PCR.

### Bioinformatics analysis

Differential gene expression was studied using R2: Genomics Analysis and Visualization Platform (http://r2.amc.nl). PCa datasets GSE50936 (Phur n=30), TCGA 2022 (n=553), GSE70768 (Dunning n=199), GSE21034 (Sawyers n=370), GSE46691 (Jenkins n=545), and GSE29079 (Sueltman n=95) were analyzed. Pairwise comparisons were performed according to data availability in each dataset: low-grade *vs* high-grade Gleason scores, benign tissue *vs* tumor tissue, and primary tumor *vs* metastasis. The analysis focused on metabolic genes sets related to steroid synthesis and steroid hormone biosynthesis, as defined by KEGG pathway annotations. CDKN1A correlation analysis in ETS fusion-negative mCRPC tumors was assessed in www.cbioportal.org (SU2C/PCF 2019).

### Statistical analyses

Gene expression data were analyzed as medians (+IQR) of fold changes relative to control, using Log2 transformation. Differential expression involved at least two experiments with three replicates, employing non-parametric Wilcoxon or Kruskal-Wallis tests for p-value calculations, followed by Dunn’s test. Proliferation and drug toxicity analyses used mean ± standard deviation with ANOVA and Dunnett’s method, considering p < 0.05 as significant. Analyses were conducted using GraphPad Prism 9.

## RESULTS

### 1. Hormone-naïve primary tumor cultures (hnPCs) exhibit molecular heterogeneity and resistance to both androgen deprivation and chemotherapy

Phenotypic and genetic characterization of ten hnPCs cultures revealed diverse tumor heterogeneity, including basal and luminal epithelial cells, neuroendocrine, and mixed neuroendocrine-epithelial phenotypes (**Figure 1A**). Cytokeratin 5 (CK5) and p63 co-expression in hnPC-05, -06, -07, -08, -09, and -10 indicated basal features, while CK5 and CK18 expression in hnPC-02 and -03 suggested mixed basal-luminal populations (**Figure 1A**). The neuroendocrine marker chromogranin A was observed in hnPC-03, -04, and -06, with hnPC-04 predominantly showing neuroendocrine features.

**Figure 1:**
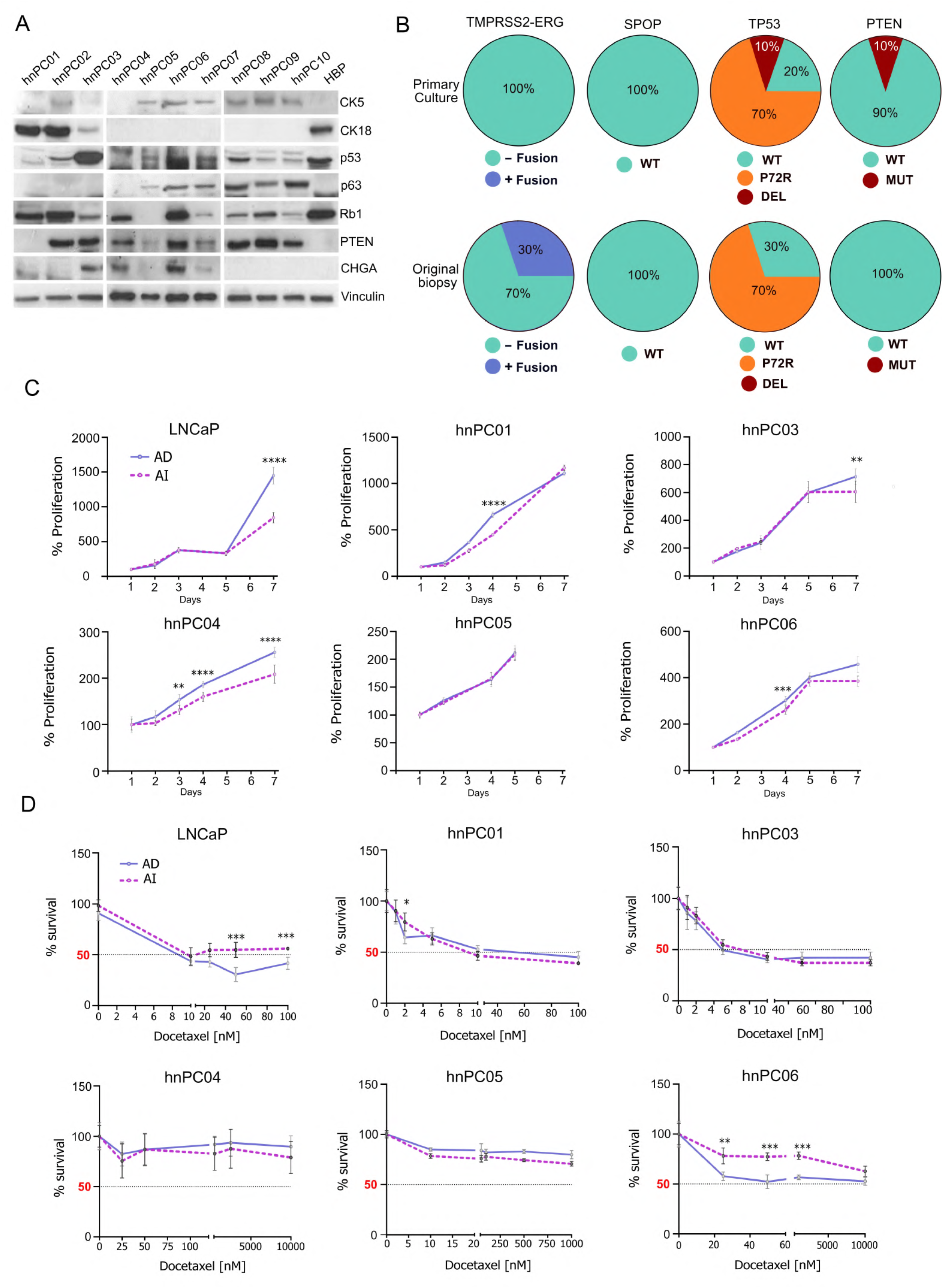
Phenotypic and genetic characterization of hnPCs cultures. **A)** <u>Phenotypic characteristics of hnPCs</u> showing expression of epithelial markers CK5, CK18 and p63, neuroendocrine marker CHGA, p53, Rb1 and PTEN expression. Vinculin expression is used as control. **B)** Gene alterations in hnPCs in comparison to original tissue biopsies (percentage). **C)** <u>Proliferation plots of LNCaP and hnPCs, AD and AI.</u> Cells plated in 96 well dishes, were grown for 7 days with (AD) or without (AI) androgens in the culture medium. Data are the mean ±S.D. of three independent experiments, each performed in octuplicate. **D)** <u>Docetaxel resistance capacity in LNCaP and hnPCs, AD and AI cells</u>. Cells were treated with vehicle (DMSO) or serial concentrations of drug for 72h. Results were from three independent experiments (performed in quintuplicate) plotted as mean ± S.D.

DNA Sequencing and protein analysis identified clinically relevant mutations in genes associated with aggressive disease progression and therapy resistance. hnPC-01 harbored a *TP53* frameshift mutation at residue 39 and a PTEN start codon mutation (M1R), correlating with loss of both p53 and PTEN protein expression (**Figure 1B, Supplementary Figure S2**). The TP53 codon 72 polymorphism (P72R, rs1042522) was present in all hnPCs except hnPC-06. Of note, the TMPRSS2–ERG fusion, reported in up to 50% of primary prostate tumors (29), was absent in all hnPCs cultures despite been detected in three original biopsy tissues, suggesting loss during culture establishment (**Supplementary Figure S3**).

To assess androgen dependence, we compared proliferation of hnPCs cultured under androgen-dependent (AD) and androgen-independent (AI) conditions with LNCaP controls (**Figure 2C**). Unlike LNCaP cells, which showed marked reduction in proliferation under androgen deprivation, most hnPCs exhibited minimal differences between AD and AI conditions, with only hnPC-04 displaying reduced growth in the absence of androgens. Particularly, hnPC-04, -05, and -06 demonstrated substantially lower proliferation rates compared to LNCaP, hnPC-01, and -03 under both conditions, suggesting that hnPC adaptation to androgen independence may involve mechanisms beyond canonical androgen signaling.

**Figure 2.**
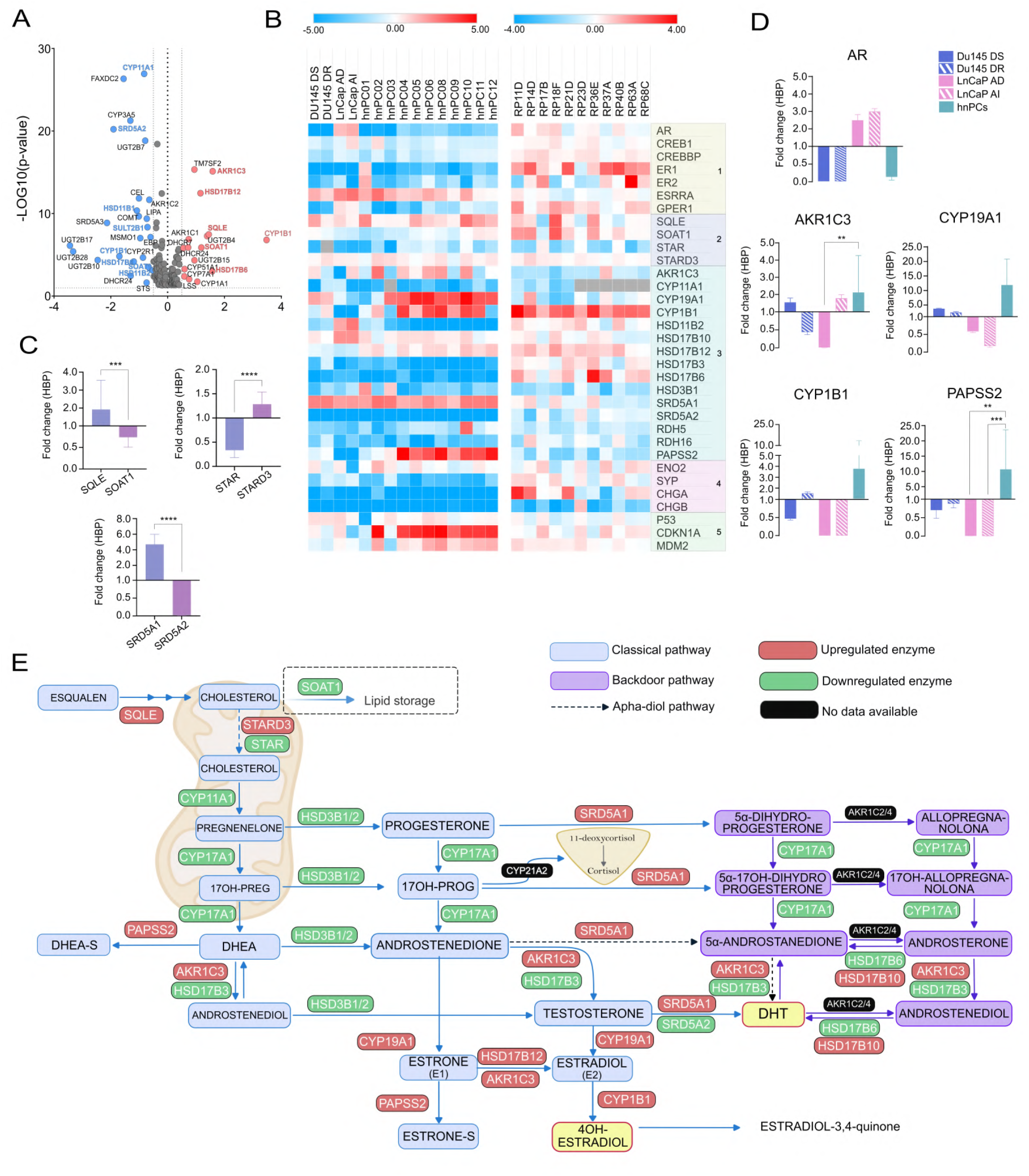
Steroidogenic pathway analysis in PCa cell lines, hnPCs and RP. **A)** <u>Volcano plot of genes of the steroidogenic pathway in PCa datasets</u>. Fold changes (Log2) and p-value (-Log10) for each gene obtained across datasets, were plotted in a volcano plot, with significance defined as p = 0.05 (−log10 p = 1.3) and fold change ≥ 1.5 (log2 FC = 0.6), indicated by dashed lines. **B)** <u>Heatmap showing expression of 35 of steroidogenic genes in PCa cell lines, hnPCs and RP</u>. Upregulated genes are shown in red, downregulated genes in blue. Molecular pathways implicated are indicated in colors. 1: Steroid Hormone Receptor signaling; 2: Steroid Metabolism; 3: Steroid hormone metabolism; 4: Sterol binding Neuroendocrine; 5: p53 signal transduction. **C)** <u>Bar plots comparing the expression of 6 steroidogenic genes in hnPCs</u>. Data represents median (+ IQR) of fold change of 11 samples in triplicates. P-values were calculated using Wilcoxon matched-pairs test. **D)** <u>Comparative expression of genes involved in the *de novo* hormone synthesis in hnPCs and PCa</u> <u>cell lines.</u> *AKR1C3,* estrogenic enzymes *CYP19A1* and *CYP1B1,* and *PAPSS2* are upregulated. Data are expressed as median (+ IQR) of fold change in triplicates; for hnPCs, the median (+IQR) of 11 samples in triplicates is shown. **E)** <u>Map of steroid synthesis pathway and enzyme expression profiles in hnPCs</u>. The pathway from squalene to estradiol is shown, with key enzymes colored red (overexpressed) or green (downregulated) according to expression levels in primary prostate cancer cultures. TBP and IPO8 were used as housekeeping genes. *P*-values are result of Kruskal-Walli’s test with Dunn’s Test comparisons. \**P*<0.05, \*\**P*<0.01, \*\*\**P*<0.001, \*\*\*\**P*<0.0001.

Analysis of mTOR pathway revealed variable responses to androgen deprivation across hnPC lines. hnPC-04 showed modest increases in protein synthesis and activation of mTOR components (phosphorylated ERK1/2, RPS6, and 4EBP1) under AI conditions (**Supplementary Figure S4**). In contrast, hnPC-05 and -06 exhibited reduced protein synthesis and decreased mTOR activity under AI conditions, consistent with their low proliferative capacity.

Androgen-independent cells (LNCaP, hnPC-04, -05, and -06) exhibited significantly greater resistance to docetaxel compared to their androgen-dependent counterparts (**Figure 1C**), with survival rates exceeding 50% under both AD and AI conditions, while hnPC-01 and - 03 showed survival rates below 50% with no significant differences between AD and AI cells (**Figure 1C**). These findings suggest that hnPC cells possess intrinsic resistance to docetaxel that is independent of androgen signaling and may be associated with their low proliferative activity.

### 2. hnPCs undergo steroidogenic reprogramming enabling *de novo* androgen synthesis

To investigate whether steroid hormone pathways rewiring contributes to cell survival and therapeutic resistance, we analyzed the expression of key steroidogenic enzymes in Du145, LNCaP, primary hnPCs, and radical prostatectomy (RP) tissue samples. We selected 33 genes for analysis based on public databases and published literature, including 7 genes related to hormone receptor signaling, 19 involved in steroid metabolism, 4 associated with neuroendocrine signaling, and 3 from the p53 pathway (**Figure 2A**).

Initial comparisons between AD and AI hnPC models revealed no major differences in gene expression (**Supplementary Figure S5a**), consistent with the limited functional divergence observed in proliferation assays. Combined with consistently low *AR* expression in both AD and AI hnPC cells, these results suggest potential insensitivity to exogenous androgens. Gene expression profiling revealed distinct patterns in hnPC-AD cells compared to RP tissues samples and established PCa cell lines (**Figure 2B and Supplementary Figure S6a-b**). Despite these differences, hnPCs retained key steroidogenic features commonly found in tumor tissues, including expression of *CYP1B1* (critical for estrogen metabolism) and *HSD17B10/12* (involved in backdoor androgen synthesis). These findings suggest that hnPCs maintain a partially conserved steroidogenic program that may contribute to cancer cell survival under androgen-depleted conditions.

Comparative analysis between hnPCs and their matched tumor biopsies (hnTBs) revealed similar expression levels of key genes involved in steroidogenesis and cell cycle regulation, including *AR, SRD5A1, CYP1B1*, *CDKN1A*, and *MDM2* **(Supplementary Figure S6c).** However, *PAPSS2* and *AKR1C3* were highly upregulated in hnPCs, likely reflecting adaptation to *in vitro* culture conditions that lack physiological androgens levels. Hormonal analysis of the culture medium confirmed this androgen-depleted environment, which closely mirror conditions in patients undergoing androgen deprivation therapy (**Supplementary Table S3).**

Beyond these conserved features, hnPCs displayed distinct metabolic adaptations consistent with steroidogenic reprogramming (**Figure 2C**). *SQLE* (involved in cholesterol synthesis) was upregulated, while *SOAT1* (cholesterol storage) was downregulated, suggesting that hnPCs prioritize cholesterol mobilization for steroidogenesis. This metabolic shift was accompanied by a change in the expression of intracellular cholesterol transporters, shifting from *StAR* to *STARD3*, which is often upregulated in cancers to support sustained steroidogenesis(30). Additionally, we observed a switch from *SRD5A2* (typically expressed in normal prostate) to *SRD5A1,* a much stronger activator of the AR consistent with preferential DHT production in cancer cells(31), These alterations distinguish hnPCs from both normal tissues and established cell lines, reflecting unique adaptation mechanisms that supports hormone synthesis and cellular survival in androgen-depleted conditions.

Compared to DU145 and LNCaP-AD cells, hnPCs displayed lower AR levels but showed coordinated upregulation of key enzymes involved in *de novo* androgen and estrogen synthesis, including *AKR1C3, CYP19A1, CYP1B1*, and *PAPSS2* (**Figure 2D**). Although some of these enzymes (*e.g., AKR1C3*) were also upregulated in LNCaP-AI cells, their coordinated expression in hnPCs suggests reactivation of intracrine steroidogenic pathways under androgen-depleted conditions. Together, these findings suggest that hnPCs possess a reprogrammed steroidogenic network featuring activation of *de novo* androgen synthesis and alternative estrogenic pathways (**Figure 2E**), which may contribute to therapy resistance in an AR-low context.

### 3. *CDKN1A* overexpression correlates with steroidogenic gene expression and regulates AKR1C3 in prostate cancer cells

*CDKN1A* (encoding p21^CIP1^) was significantly overexpressed in hnPC cultures compared to PCa cell lines (Kruskal-Wallis’ test, p=0.011, **Figure 3A**). The only exceptions were hnPC-01 and hnPC-03, which also showed higher proliferation rates than other hnPCs. Although p21^CIP1^ has been classically associated with cell cycle arrest and senescence, its overexpression has paradoxically been linked to aggressive tumor behavior in several cancers (32–34). Consistent with our findings, analysis of the Sawyers dataset (GSE21034, *n*=370) revealed significant *CDKN1A* upregulation in prostate tumors compared to adjacent benign tissue (ANOVA, *p* =0.016) (**Figure 3B**).

**Figure 3.**
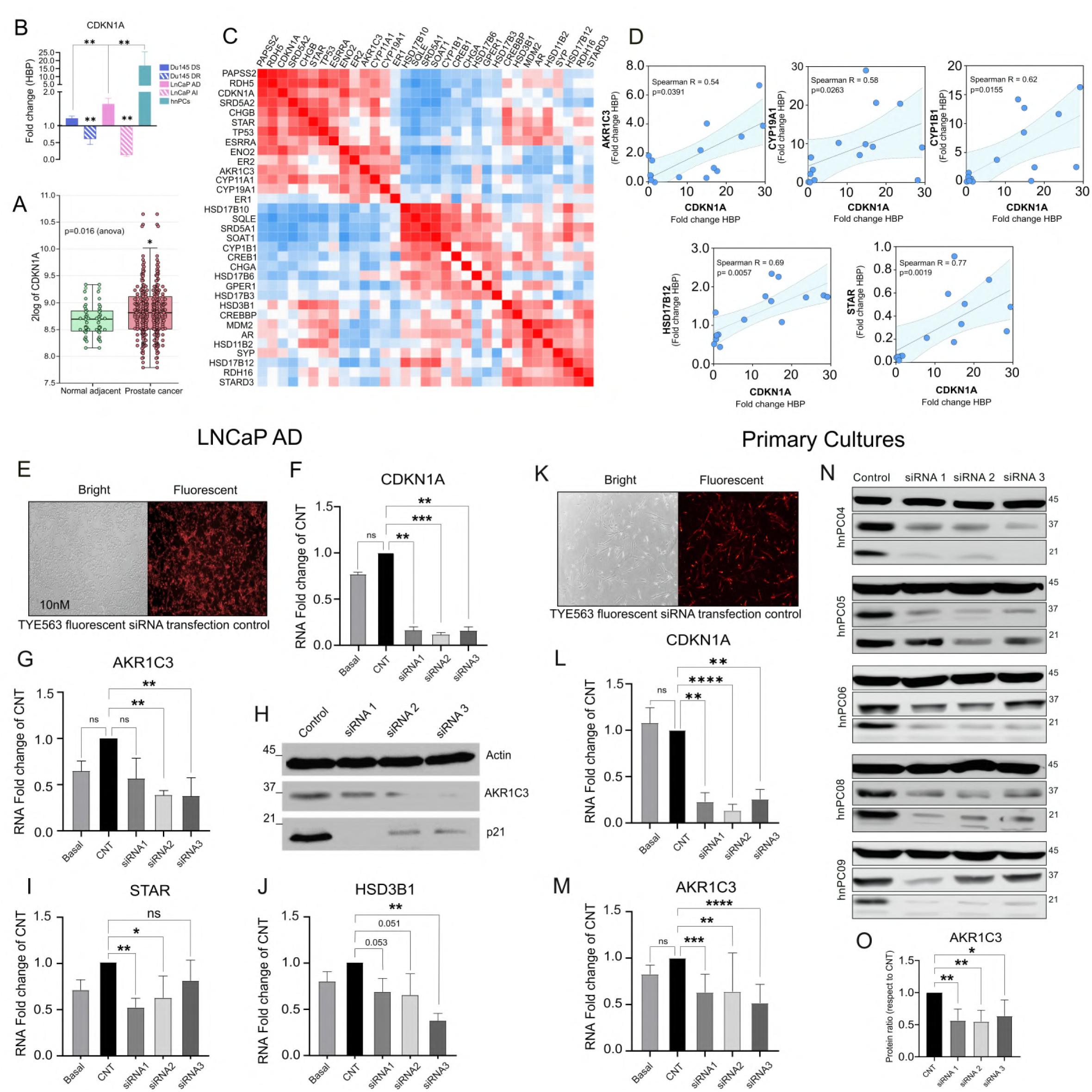
CDKN1A (p21) overexpression correlates with steroidogenic genes expression. A-B) <u>CDKN1A expression in PCa tissues and hnPCs</u>**. A:** Box plot representing CDKN1A expression in PCa tissues (n=300) in comparison with normal adjacent tissue (n=70) from Sawyers dataset using GenomicR2. Significant difference assessed by Anova (p<0.016). **B:** CDKN1A expression in PCa cell lines and hnPCs (in triplicates), presented as median +IQR. **C-D)** <u>Correlations between CDKN1A and steroidogenic genes</u>. **C:** Multiple Spearman correlation matrix highlighting clustering of CDKN1A with several steroidogenic genes. **D:** Pairwise Spearman correlation analysis between CDKN1A and selected steroidogenic genes. **E-F)** <u>Knockdown of p21 in LNCaP cells</u>. Transfection efficiency is shown by fluorescence imaging (bright-field and red), and RNA analysis confirms p21 knockdown. **G-J)** p21 knockdown in LNCaP is associated with downregulation of AKR1C3 RNA and protein (Western blot), as well as decreased expression of STAR (**I**) and HSD3B1 (**J**). **K-L)** <u>Knockdown of p21 in hnPCs.</u> Transfection efficiency is shown by fluorescence, and RNA analysis confirms p21 inhibition. **M-O**) p21 knockdown in hnPCs is associated with AKR1C3 RNA downregulation, corroborated by Western blotting of five independent cultures (**M–P**) and quantified by protein densitometry (**O**). Protein levels were normalized to β-actin and compared to control cultures. All experiments were performed in triplicate in at least two independent assays. RNA data presented are median + IQR. *P*-values were obtained by Kruskal-Walli’s test with Dunn’s multiple comparisons. Protein quantification for AKR1C3 was obtained by scanning densitometry of the band area, normalized to β-actin. **ns:** no significant *P<0.05, \*\**P*<0.01, \*\*\**P*<0.001, \*\*\*\**P*<0.0001

To determine whether *CDKN1A* upregulation is associated with steroidogenic gene activity, we performed Spearman’s correlation analysis in hnPC cells. *CDKN1A* positively correlated with multiple key steroidogenic genes, including *STAR, HSD17B12, SOAT, SRD5A1, CYP1B1,* and especially *AKR1C3* (R>0.5, p-value < 0.05; **Figure 3C-D**). These associations suggested that CDKN1A may modulate components of the steroidogenic pathway.

To test this hypothesis, *CDKN1A* was knocked down in LNCaP-AD cells and hnPCs using three independent siRNAs. Knockdown efficiency exceeded 80% in all cases and consistently led to reduced *AKR1C3* mRNA levels across both cell types (**Figures 3E-F, 3K-L**). The effects on *STAR* and *HSD3B1* were variable (**Figure 3I-J)**. Western blotting analyses further confirmed a significant decrease in AKR1C3 protein levels in p21-knockdown LNCaP cells and in hnPC-04, -05 and -08 cultures (**Figures 3G-H, 3M-O**).

Together, these data indicate that *CDKN1A* may regulate steroidogenic pathway activation in prostate cancer, in part by promoting expression of key enzymes such as AKR1C3. The consistent downregulation of AKR1C3 following p21^CIP1^ knockdown supports a mechanistic link between p21 overexpression and enhanced intratumoral *de novo* androgen synthesis in hnPC models.

### 4. Distinct p21 subcellular localization in LNCaP cells and hnPCs suggests distinct functional roles, with hnPCs exhibiting features of partial senescence

The functional activity of p21^CIP1^ is strongly influenced by its subcellular localization: nuclear p21 typically mediates tumor-suppressor cell-cycle arrest, whereas cytoplasmic p21 is associated with pro-survival and oncogenic functions (35). Understanding the determinants of its localization is therefore essential for interpreting its role in prostate cell biology.

In LNCaP-AD cells, subcellular fractionation showed p21 predominantly in the cytoplasmic fraction (**Figure 4A**). Immunofluorescence confirmed this pattern, revealing strong perinuclear and cytoplasmic staining in most cells, with only a minority displaying exclusive nuclear localization (**Figure 4B**, white arrow). This primarily cytoplasmic distribution suggests a potential that p21 in LNCaP-AD cells, may engage in non-canonical, potentially pro-oncogenic roles.

**Figure 4.**
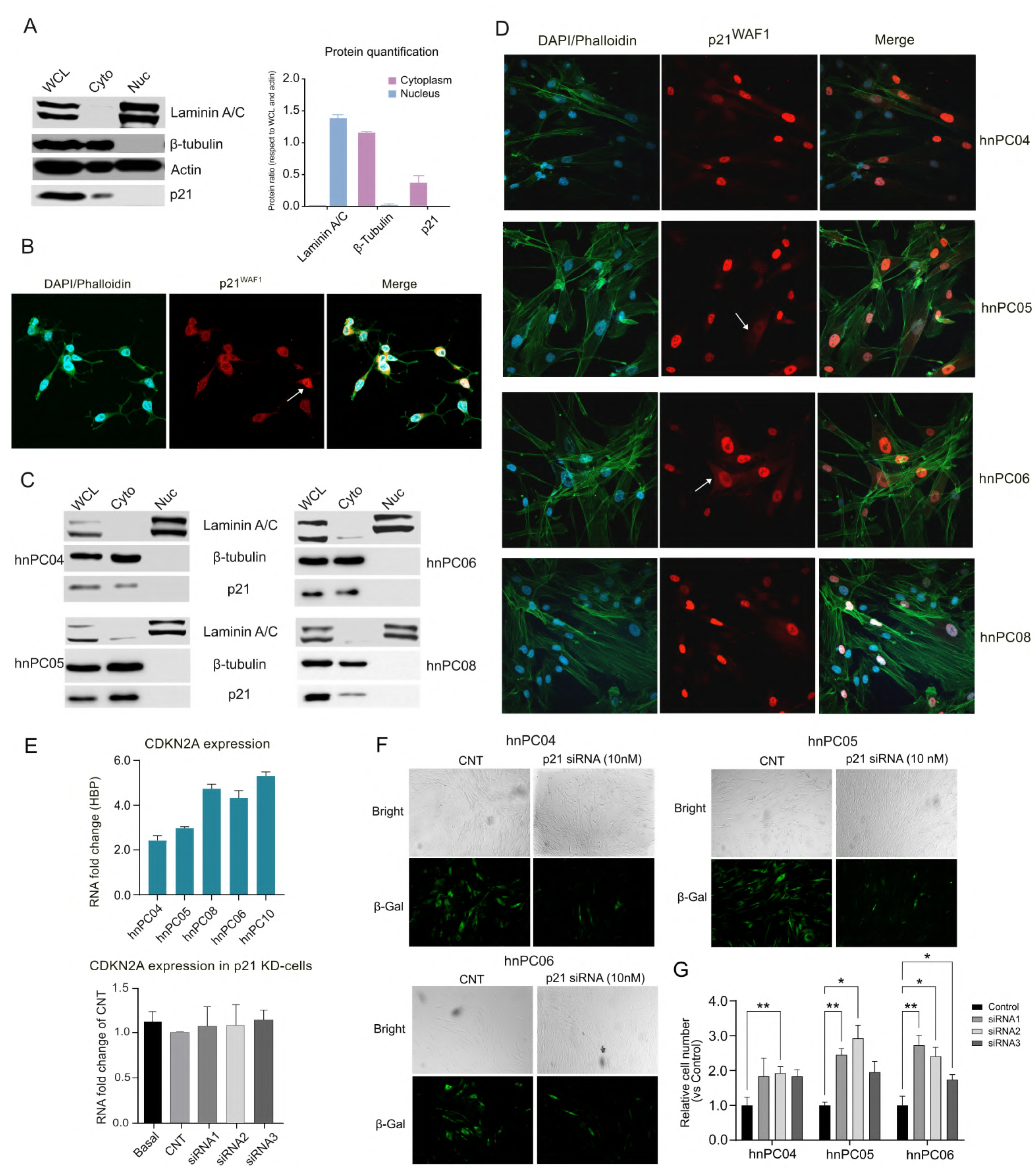
Senescent characteristics of hnPCs. **A)** <u>Subcellular localization of p21 in LNCaP cells.</u> Representative western blot of whole cell lysate (WCL), cytoplasmic (Cyto), and nuclear (Nuc) fractions, showing p21 is predominantly observed in the cytoplasm. Lamin A/C (70–62 kDa) and β-tubulin (55 kDa) were used as nuclear and cytoplasmic controls, respectively, confirming correct fractionation. Protein bands (triplicates) were quantified and normalized to β-actin and compared to WCL (ratio = 1). **B)** <u>Immunofluorescence of p21 in LNCaP cells.</u> Confocal microscopy images showing p21 (Alexa-594, red), nucleus (DAPI, blue), and actin (Phalloidin, green). p21 is primarily detected perinuclear and cytoplasm. Minor in nuclear localization in white arrow (25X magnification). **C)** <u>Subcellular localization of p21 in hnPCs</u>. Western blot analysis of nuclear and cytoplasmic fractions shows p21 primarily in cytoplasm. Lamin A/C and β-tubulin were used as nuclear and cytoplasmic controls, respectively, confirming correct fractionation. **D)** <u>Immunofluorescence of p21 in hnPCs</u>. Primary cultures stained for p21 (Alexa-594 red), nucleus (DAPI-Blue) and actin (Phalloidin-Green). p21 is more frequently detected in thr nucleus. Cells showing both nuclear and cytoplasmic p21 are indicated in white arrows). (25X magnification). **E)** <u>Senescence marker CDKN2A in hnPCs</u>. Cells express CDKN2A mRNA (Log2 fold change relative to BPH), confirming senescent state. Knockdown of p21 does not alter CDKN2A expression. **F-G)** <u>Senescence-associated β-galactosidase activity and proliferation after p21 knockdown</u>. hnPCs were transfected with control siRNA or p21-targeting siRNAs (#1–3) for 48 h after 24 h of growth. **F**: Brightfield images show total cell density, fluorescence images show B-gal activity. In these panels, siRNA #2 is shown as a representative knockdown. Control cells exhibit high B-gal staining, whereas p21 knockdown cultures show fewer B-gal-positive cells and increased cell density. **G**: Quantification of relative cell number for all three siRNAs across three cultures is shown (mean ± SD from five random fields per well). All experiments were performed in triplicate.

In contrast, hnPC cultures displayed a markedly different pattern. Although p21 was again detected exclusively in the cytoplasmic fraction by biochemical analysis (**Figure 4D**), immunofluorescence revealed predominantly nuclear staining, with only occasional cytoplasmic signal (white arrow). This discrepancy likely reflects methodological differences in sensitivity and fraction purity, emphasizing the need for complementary approaches when assessing protein localization. The nuclear enrichment observed by microscopy suggests that in hnPCs, p21 may primarily contribute to growth arrest and senescence-associated functions.

To further characterize senescence in hnPCs, we examined *CDKN2A* (p16^INK4a^) expression (**Figure 4E**) and senescence-associated β-galactosidase (SA-β-gal) activity (**Figure 4F**). *CDKN2A* expression was upregulated in hnPC-04, -05, -06, and -10, and its expression was unaffected by p21-knockdown. In contrast, strong SA-β-gal activity was observed in control hnPCs but was markedly reduced following p21 knockdown, accompanied by increased cell density indicative of enhanced proliferation (**Figure 4G**). Thus, p21^CIP^ is required to maintain SA-β-gal-associated senescence features, whereas CDKN2A induction alone is insufficient to sustain the senescence phenotype in the absence of p21. These observations indicate that p21 and CDKN2A/p16 act through parallel, complementary pathways to regulate senescence and cell-cycle arrest.

Senescence is also characterized by SASP-associated signaling, which are regulated by stress-responsive signaling pathways. Given the role of Notch1 in early senescence and SASP modulation, we assessed Notch1 levels following CDKN1A knockdown. Notch1 levels increased in hnPC-04 and -06, whereas only showed modest changes in -05, and remained unchanged in -08 (**Supplementary Figure S7**), This heterogeneity suggests that p21 loss does not uniformly abrogate senescence-associated signaling but instead triggers context-dependent remodeling of Notch1-associated pathways in a subset of hnPCs.

### 5. *CDKN1A* expression correlates with steroidogenic genes in ETS fusion-negative mCRPC tumors naïve to AR signaling inhibitors

To assess whether the observed *CDKN1A* expression patterns *in vitro* reflect regulatory programs relevant to endogenous steroidogenesis and lineage plasticity, we interrogated publicly available transcriptomic datasets selecting ETS-fusion-negative metastatic CRPC tumors from patients who did not receive androgen receptor signaling inhibitors (ARSIs) (36). *CDKN1A* expression was moderately higher in ETS fusion-negative tumors (median = 4.811; mean = 4.689) compared to ETS fusion-positive tumors (median = 4.396; mean = 4.495), although difference did not reach statistical significance (*p = 0.088*) (**Figure 5A**).

**Figure 5.**
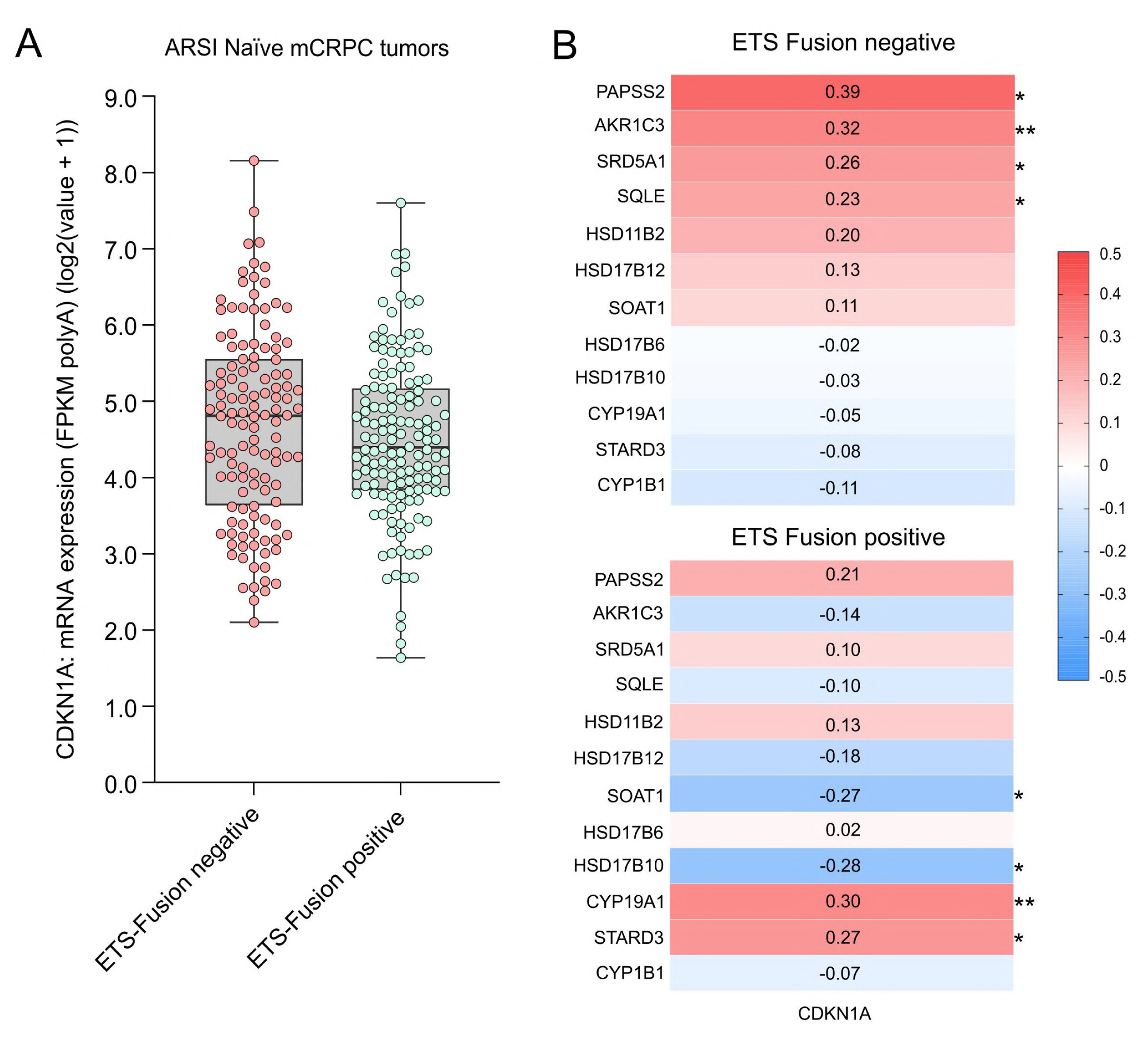
CDKN1A expression correlates with steroidogenic genes in ETS fusion-negative mCRPC tumors. **A)** CDKN1A expression in ETS fusion-positive and fusion-negative tumors. Median values were higher in fusion-negative samples, although not statistically significant (*p* = 0.088). **B)** Correlation matrices showing the association between CDKN1A and steroidogenic genes in ETS fusion-negative (top) and fusion-positive (bottom) tumors. Stronger and more consistent correlations were observed in fusion-negative tumors. Data was obtained from publicly available mCRPC transcriptomic datasets (Abida et al. 2019). Correlations were computed using Spearman’s rank correlation coefficient.

Next, we examined whether *CDKN1A* expression was associated with genes involved in steroidogenesis. In ETS fusion-negative tumors, *CDKN1A* showed positive significant correlations with several key steroidogenic enzymes, including *AKR1C3* (r = 0.32, *p < 0.01*), *PAPSS2* (r = 0.39, *p < 0.05*), *SRD5A1* (r = 0.26, *p < 0.05*), and *SQLE* (r = 0.23, *p < 0.05*). In contrast, ETS-fusion positive tumors exhibited weaker and more variable correlations. In this group, *CDKN1A* was negatively correlated with *AKR1C3* (r= –0.14), while showing positive correlation with *CYP19A1* (r = 0.30, *p* < 0.01) (**Figure 5B**).

These findings suggest that in ETS-negative, ARSI-naïve tumors, elevated *CDKN1A* expression may contribute to enhanced steroidogenic signaling. Taken together, these data suggest a link between senescence-associated pathways and steroidogenic reprogramming in this tumor subset, in line with our *in vitro* observations.

## DISCUSSION

This study establishes hormone-naïve primary tumor cultures (hnPCs) as valuable preclinical models that recapitulate key features of aggressive prostate cancer, including therapeutic resistance and metabolic reprogramming. Our findings reveal an unexpected link between *CDKN1A* overexpression, partial senescence, and coordinated upregulation of steroidogenic enzymes, particularly AKR1C3. This molecular signature supports intratumoral androgen synthesis, conferring resistance to ADT and chemotherapy, providing insight into mechanisms driving early progression to castration-resistant prostate cancer.

Primary cultures preserved molecular diversity and phenotypic hallmarks of original tumors, consistent with prior reports (37–40), including steroidogenic enzymes expression and cell cycle regulators after *ex vivo* adaptation. Retention of basal, luminal, neuroendocrine, and mixed phenotypes, underscores tumor heterogeneity and distinguishes them from traditional homogeneous traditional cell lines (22,23).

The loss of *TMPRSS2-ERG* fusion in three tumor biopsies during cultures establishment resulted in enrichment of ETS-negative cells, which enhance the clinically relevance for studying ARSI-naïve, ETS-negative mCRPC, an underexplored subset with distinct therapeutic vulnerabilities (41). The intrinsic resistance to ADT and docetaxel, combined with low proliferation, suggests hnPCs exist in a quiescent state refractory to conventional therapies, aligning with clinical observations that slow-cycling populations serve as reservoirs for resistance and recurrence (42,43).

Our data demonstrates a coordinated steroidogenic reprogramming characterized by upregulation of enzymes involved in *de novo* androgen and estrogen synthesis. The shift from *STAR* to *STARD3* for cholesterol transport, coupled with increased *SQLE* and decreased *SOAT1*, indicates metabolic prioritization of cholesterol mobilization for steroid biosynthesis (44). While StAR is hormonally regulated and expressed mainly in classical steroidogenic tissues, *STARD3* is frequently upregulated in breast and prostate cancer where it allows sustained steroidogenesis, energy and rapid growth (45). The switch from *SRD5A2* to *SRD5A1* reflects adaptation toward pathways preferentially utilized in cancer for DHT production, such as the backdoor and 5α-dione involving HSD17B6/10/12 (46,47). The marked overexpression of AKR1C3, which converts weak androgens to testosterone, is significant, given its established role as a key mediator of castration resistance by enabling tumors to synthesize androgens from adrenal precursors, sustaining AR signaling despite systemic suppression(48,49). However, in our models, the steroidogenic signature observed emerges in the context of reduced AR expression, highlighting a potential AR-independent mechanism of adaptation to maintain hormonal support for growth and survival through alternative, intracrine pathways (50). Consistent with this notion, *CYP19A1* (aromatase) upregulation, also reinforce the concept that hnPCs engage alternative steroidogenic routes beyond canonical AR signaling through estrogen synthesis, previously shown to promote tumor cell survival and progression in prostate cancer under androgen-deprived conditions (51,52)

The overexpression of *CDKN1A* in hnPCs, particularly in cultures with low proliferative capacity, challenges the traditional view of p21^CIP1^ solely as a tumor suppressor. Our findings suggest that, in aggressive prostate cancer, p21^CIP1^ may acquire non-canonical functions promoting tumor progression (53–55). Strong positive correlations between *CDKN1A* and steroidogenic enzymes expression, coupled with consistent AKR1C3 downregulation following p21 knockdown, support a mechanistic link between p21 and metabolic reprogramming. In contrast, AR-positive LNCaP cells displayed negative correlation between CDKN1A and AKR1C3 under both androgen-dependent and -independent conditions, consistent with preserved AR signaling. In this context, AR activation may be repressing *CDKN1A* expression (56,57) suggesting that canonical AR activity constrains the p21-driven steroidogenic program observed in hnPCs. This contrast indicates that the pro-tumorigenic functions of p21^CIP1^ are not intrinsic but emerge specifically in AR-deficient contexts, highlighting the context-dependent role of p21.

The mechanism by which p21 regulates AKR1C3 remains to be fully elucidated. Possibilities may include p21 interaction with transcription factors regulating steroidogenic gene promoters such as NF-kB (58), influence on chromatin remodeling (59) favoring steroidogenic gene expression, or p21-mediated cell cycle arrest indirectly promoting metabolic reprogramming (60,61) by shifting resources from proliferation to steroid biosynthesis.

The cytoplasmic p21 localization in LNCaP cells aligns with reports associating cytoplasmic p21 with oncogenic functions, including survival promotion and therapy resistance in different cancers (62–64). In hnPCs, the discrepancy between fractionation analysis (cytoplasmic p21) and immunofluorescence (nuclear signal) requires careful interpretation. Biochemical fractionation can be influenced by nuclear membrane integrity and centrifugation parameters, low temperatures, potentially causing artificial redistribution (65,66). Additionally, p21 shuttles between nucleus and cytoplasm in response to stress(67,68). Immunofluorescence data, preserving spatial relationships in intact cells, likely provides more accurate representation.

Co-expression of nuclear p21, p16, and SA-β-galactosidase activity alongside limited proliferative capacity suggests hnPCs exist in a partial senescence state (69). Functional assays indicate that p21 is required to maintain specific senescence-associated features, In contrast, the persistence of p16 overexpression following p21 knockdown supports the existence of parallel regulatory pathways, in which p21 sustains selected senescence traits while simultaneously enabling plasticity, facilitating non-canonical functions and tumor progression (33,70). Notably, increased proliferation following p21 depletion despite sustained p16 overexpression indicates that p16 alone is insufficient to enforce stable cell-cycle arrest and may exert context-dependent pro-proliferative functions, as reported in some cancer models (71,72).

Notch1, a key regulator of SASP architecture reinforcing juxtacrine signaling, has been shown to exert context-dependent tumor-suppressive or tumor-promoting effects in various cancer types (73–75). Heterogeneous Notch1 responses observed upon p21 depletion in a subset of hnPCs are consistent with its dual functionality and align with the increased proliferation observed in that subset of cultures. Although functional validation will be required, these findings underscore that p21 inhibition alone does not uniformly abrogate tumor-promoting properties, highlighting the resilience of partial senescence programs and the need to consider combinatorial targeting strategies.

Interrogation of independent transcriptomic datasets strengthens the translational relevance of our findings. Positive correlations between CDKN1A and steroidogenic enzymes, specifically in ETS-negative, ARSI-naïve mCRPC suggest this regulatory network operates in a defined molecular subset, while absence in ETS-fusion-positive tumors indicates regulation by different molecular drivers (76,77). These findings enable patient stratification. ETS-negative tumors (approximately 50% of prostate cancers) may be particularly susceptible to therapies targeting steroidogenic enzymes, when CDKN1A is elevated. The enrichment of this signature in ARSI-naïve patients suggests it may represent an adaptive response predisposing tumors to castration resistance through pathways such as SPINK1 signaling (78,79). CDKN1A expression, potentially combined with AKR1C3, could serve as a predictive biomarker identifying high-risk patients who might benefit from aggressive upfront combination therapies.

Our findings reveal potential therapeutic strategies. First, targeting AKR1C3 in combination with ADT may be effective in patients with elevated CDKN1A. Second, disrupting partial senescence, either promoting full senescence or reversing it, warrants exploration. Third, understanding how p21 subcellular localization modulates its oncogenic versus tumor-suppressive functions may identify strategies to restore tumor suppression. This senescence-steroidogenesis connection highlights the context-dependent role in cancer. While serving as a barrier in early tumorigenesis, our data suggest it may support tumor survival later, underscoring caution in senescence-targeting therapies.

## CONCLUSIONS

These findings reveal a previously underrecognized connection between CDKN1A/p21^CIP1^ overexpression, partial senescence, and steroidogenic reprogramming in aggressive, hormone-naïve, ETS-negative prostate cancer models. hnPCs display a senescence-like phenotype marked by p21^CIP1^, p16^INK4a^, and SA-β-galactosidase, along with sustained low-proliferation rates, suggesting senescence bypass with acquisition of oncogenic properties rather than irreversible growth arrest. The functional link demonstrated by CDKN1A knockdown, regulating AKR1C3 expression, indicates a novel regulatory axis driving intratumoral androgen and estrogen synthesis, which may contribute to therapy resistance and tumor progression. These observations provide mechanistic insight into how senescence-associated pathways can paradoxically support adaptation, low-level proliferation, and intracrine steroidogenesis. Validation in ETS-negative, ARSI-naïve mCRPC clinical samples suggests that this molecular signature could serve as a predictive biomarker and guide the development of targeted therapeutic strategies to disrupt this adaptive program early in disease, prior to the emergence of full castration resistance.

## Limitations

Despite providing novel insights into the interplay between CDKN1A/p21^CIP1^, partial senescence, and steroidogenic features, several limitations should be acknowledged. First, while hnPCs retain key molecular features, they lack the complex tumor microenvironment that influences behavior and therapeutic responses *in vivo*. Second, the sample size of twelve hnPC models may not fully capture the molecular heterogeneity of ETS-negative prostate cancers. Third, although correlations between p21 and AKR1C3, as well as other steroidogenic genes, were observed, functional validation *in vivo* is lacking, and mechanistic pathways remain to be fully elucidated. Fourth, the nuclear versus cytoplasmic duality of p21 was primarily assessed *in vitro*, and its relevance in tumor tissues requires confirmation. Finally, transcriptomic analyses in ETS-negative, ARSI-naïve mCRPC cohorts are retrospective and correlative, limiting causal inference. Future studies addressing these limitations will be crucial to validate the translational relevance of the senescence–steroidogenesis axis and its potential as a therapeutic target.

## Data availability

The data that support the findings of this study are available from the corresponding author upon reasonable request.

## Ethics statement

This study was approved by the Medical Research Ethics Committee of Vall d’Hebron Hospital (protocol number: PR(AG) 96/2015). The studies were conducted in accordance with the local legislation and institutional requirements. The participants provided their written informed consent to participate in this study.

## Acknowledgments

The authors extend their gratitude to the Biochemistry service from Hospital Vall d’Hebron, Barcelona-Spain, for technical support.

## Author contributions

E.T.G. and R.P. conceived and designed the study.

E.T.G., V.M., and J.A.M. developed the methodology.

E.T.G., V.M., J.A.M., and J.G. performed experiments and data analysis.

J.P., A.C., B.M., M.M., M.E.S., and R.S. contributed to resources and data/sample acquisition.

E.T.G. wrote the original draft and prepared the figures.

R.P., T.T., R.F., and J.M. critically revised the manuscript.

R.P. supervised the study and acquired funding.

**The authors declare no conflicts of interest.**

