## Supplementary Material for "p21^CIP1/WAF1^-mediated partial senescence supports steroidogenesis and therapy resistance in primary prostate cancer cultures"

**Table S1. Patient's characteristics**

| | **At diagnosis** | **At study completion** |
|---|---|---|
| **Age, median (IQR)** | 76 (64-85) | |
| **PSA, median (IQR)** | 87.75 (39-222) | 0.74 (0.3-60.62) |
| **Gleason grade, median (min-max)** | 8.5 (8-10) | - |
| **Metastatic event, n (%)** | 1 (6.25%) | 13 (81.25%) |
| **CRPC event, n (%)** | 0 (0%) | 9 (56.25%) |
| **Exitus event, n (%)** | 0 (0%) | 10 (62.5%) |
| **Exitus (other causes)** | - | 3 (18.75%) |
| **Exitus (PCa)** | | 7 (43.75) |

**Table S2**. TaqMan assays for RT-qPCR

| GENE | Catalog number | TaqMan assay | Amplicon (bp) |
|---|---|---|---|
| **AR** | 4453320 | Hs00171172_m1 | 72 |
| **ESR1** | 4453320 | Hs01046816_m1 | 65 |
| **ESR2** | 4453320 | Hs01100353_m1 | 73 |
| **ESRRA** | 4453320 | Hs01067166_g1 | 146 |
| **GPER1** | 4453320 | Hs01922715_s1 | 136 |
| **SIRT1** | 4453320 | Hs01009006_m1 | 91 |
| **STAR** | 4448892 | Hs00986559_g1 | 78 |
| **STARD3** | 4448892 | Hs00199052_m1 | 90 |
| **AKR1C3** | 4453320 | Hs00366267_m1 | 112 |
| **CYP1B1** | 4453320 | Hs00164383_m1 | 118 |
| **CYP19A1** | 4453320 | Hs00903411_m1 | 72 |
| **HSD3B1** | 4448892 | Hs04194787_g1 | 112 |
| **HSD17B3** | 4448892 | Hs00970004_m1 | 125 |
| **HSD17B10** | 4448892 | Hs00189576_m1 | 76 |
| **HSD17B12** | 4448892 | Hs00275054_m1 | 65 |
| **SRD5A1** | 4448892 | Hs00971645_g1 | 100 |
| **SRD5A2** | 4448892 | Hs00936406_m1 | 105 |
| **PAPSS2** | 4331182 | Hs00989928_m1 | 69 |
| **ENO2** | 4453320 | Hs00157360_m1 | 77 |
| **SYP** | 4453320 | Hs00300531_m1 | 63 |
| **CHGA** | 4453320 | Hs00900370_m1 | 67 |
| **CHGB** | 4453320 | Hs01084631_m1 | 112 |
| **TP53** | 4453320 | Hs01034249_m1 | 108 |
| **MDM2** | 4453320 | Hs00540450_s1 | 104 |
| **CDKN1A** | 4453320 | Hs00355782_m1 | 66 |
| **CDKN2A** | 4453320 | Hs00923894_m1 | 115 |
| **TBP** | 4453320 | Hs00427620_m1 | 91 |
| **IPO8** | 4453320 | Hs00914057_m1 | 88 |

**Table S3.** Biochemical analysis of hormones (analytes) content in serum and culture medium

| Analyte | Concentration in FBS | Concentration in medium | Reference values in blood* |
|---|---|---|---|
| Cholesterol (mg/dL) | 34.0 | 2.38 ↓ | 50-200 |
| Triglycerides (mg/dL) | 65.0 | 4.55 ↓ | 35-150 |
| Estradiol (pg/mL) | 11.0 | 0.77 ↓ | 20-350 |
| SHBG (nmol/L) | 1.5 | 0.11 ↓ | 23-160 |
| Testosterone total (ng/dL) | 5.0 | 0.35 ↓ | 300-1000 |
| Cortisol (µg/dL) | 0.64 | 0.045 ↓ | 5.27-22.5 |
| DHEA-S (µg/dL) | 2.0 | 0.14 ↓ | 100-375 |
| DHEA (ng/dL) | 0.4 | 0.028 < | 0.06-0.7 |
| 17OH-Progesterone (ng/dL) | 0.14 | 0.0098 ↓ | 0.35-3.0 |
| Androstenedione (ng/dL) | 0.2 | 0.014 ↓ | 0.5-3.5 |
| IGF-1 (ng/dL) | 113 | 7.91 ↓ | 180-500 |
| 17OH-Pregnenolone (ng/dL) | 2.97 | 0.21 < | 0.55-5.0 |
| Estrone (pg/mL) | 270 | 18.9 = | 15-77 |

*Normal values in males from kit manufactures and Mayo Clinic (https://endocrinology.testcatalog.org/)

Concentration respect to normal values: ↓ very low, < low , = Equal

**Figure S1**. **Representative images of ten hormone-naïve tumor cultures.** Images were taken at 10× magnification under bright-field microscopy. Scale bar = 100 µm. Each image represents an independent culture.

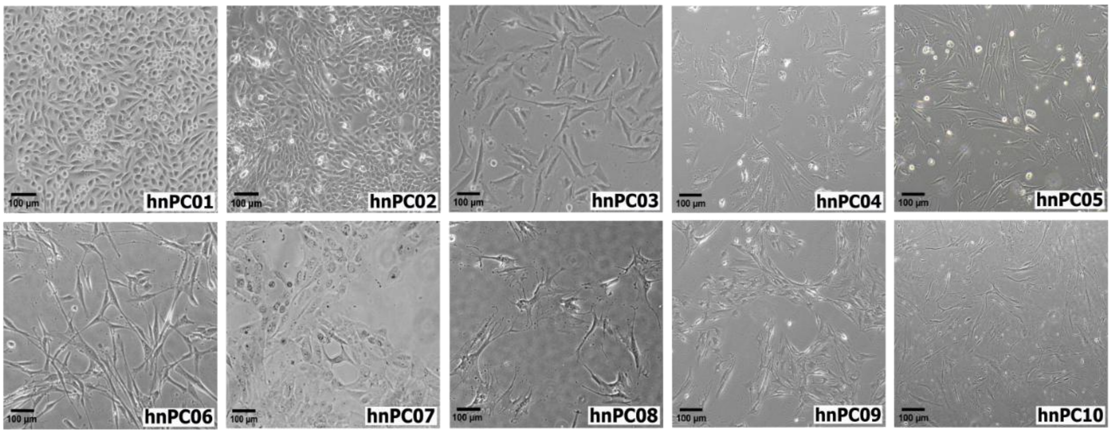


**Figure S2**. **TP53 Deletion in hnPC01 culture. a)** Reference gene (ENST00000269305)**.** showing the c.414C position (green box), corresponding to the nucleotide that is deleted in the mutant sequence, and the lysine (K) residue at position 139 in the normal open reading frame (blue arrow). **b)** Gene with a single-nucleotide deletion at c.414C. The deletion causes a frameshift mutation, resulting in an amino acid substitution from lysine (K) to arginine (R) at residue 139 (orange box) and ultimately leading to a premature stop codon at residue 169.

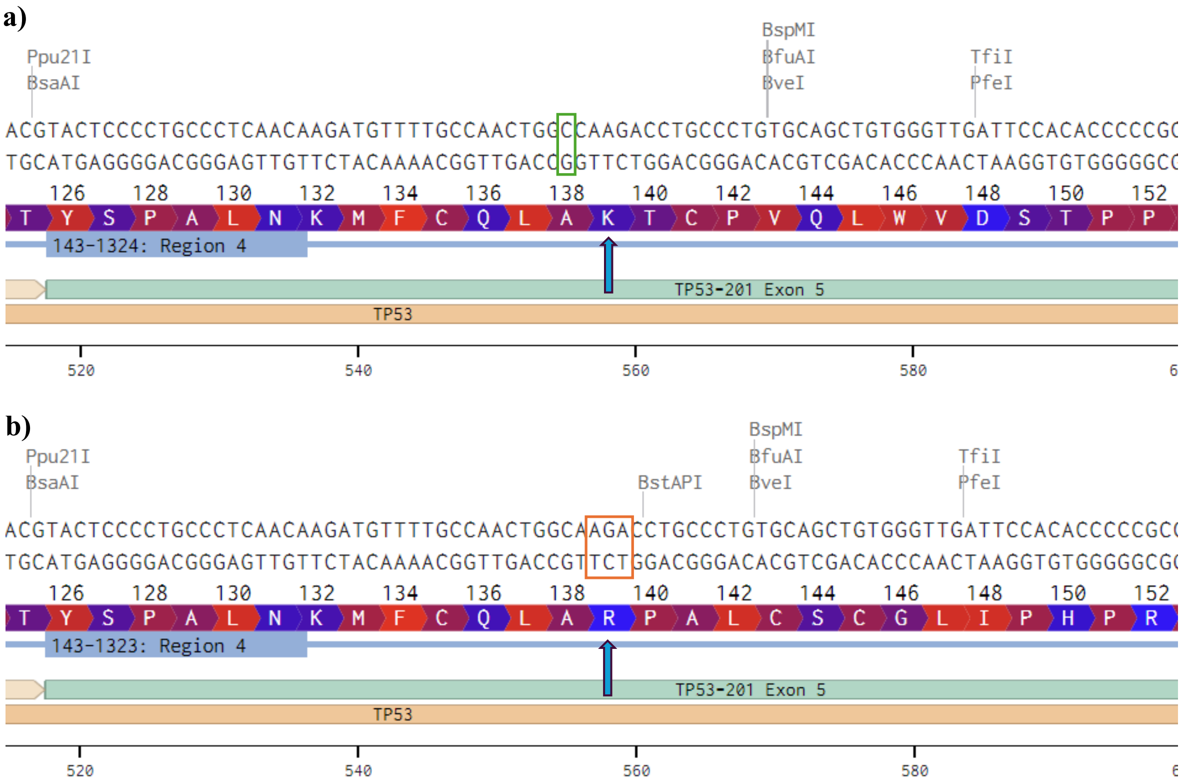

**Figure S3**. **PCR detection of TMPRSS2-ERG fusion in tumor biopsies and derived culture.** *Top:* PCR performed on 10 primary prostate tumors (PCa) using primers specific for the TMPRSS2-ERG fusion. Bands around 500–600 bp correspond to the expected fusion products. *Bottom:* Two gels run separately and digitally assembled to display all culture samples. *First gel:* molecular weight ladder (Ladder), followed by PCR controls for Rb (RbI and RbII) and PTEN (PTEN I), showing bands of approximately 800 bp. Subsequently, tumor-derived cultures 1–5 are shown. *Second gel:* tumor-derived cultures 6–10, followed by TMPRSS2 PCR controls (~500–600 bp).

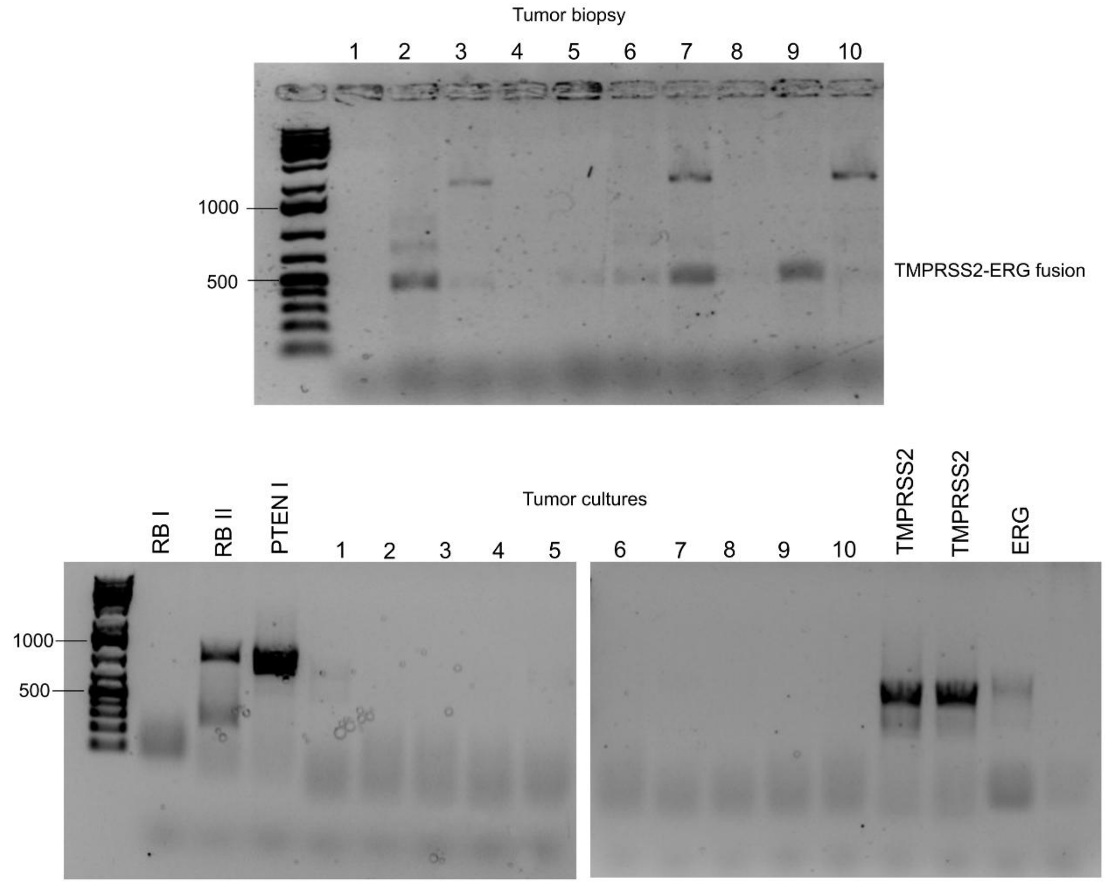

**Figure S4. Characterization of protein synthesis and gene expression in hormone-naïve tumor cultures. a)** Representative immunoblots showing protein synthesis ratios in LNCaP AD and AI, Du145 DS and DR, and hnPCs treated with 10 µg/mL puromycin and 10 µg/mL cycloheximide. Negative controls are untreated cells**. b)** Analysis of the mTORC1 pathway. Shown are levels of the indicated proteins in AD and AI LNCaP and Du145 DS and DR cells (left panel) and hnPCs (right panel). Minor differences are observed between AD and AI.

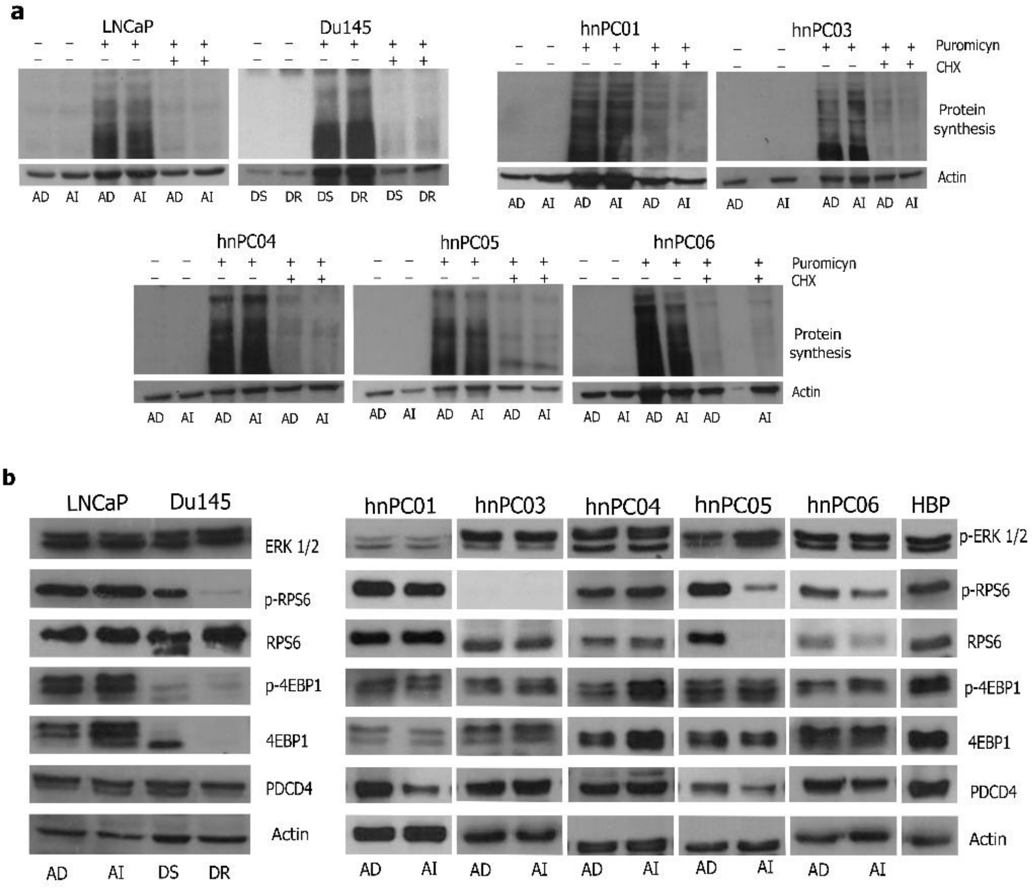

**Figure S5.** Comparison between AD and AI models for the expression of steroidogenic genes and self-renewal markers. **a)** Exploratory characterization of primary cultures. Heatmap of gene expression in three representative hnPCs. Data represents mean fold changes relative to control (BPH) in triplicate. AR is underexpressed in both AD and AI, and most genes show similar expression levels, supporting the decision to focus on AD cultures. **b)**. Relative mRNA levels were with respect to AD models. TBP and IPO8 were used as housekeeping genes. Data are the mean ± S.D of two experiments, each performed in sextuplicate. * P<0.05, **P<0.01, ***P<0.001, ****P<0.0001

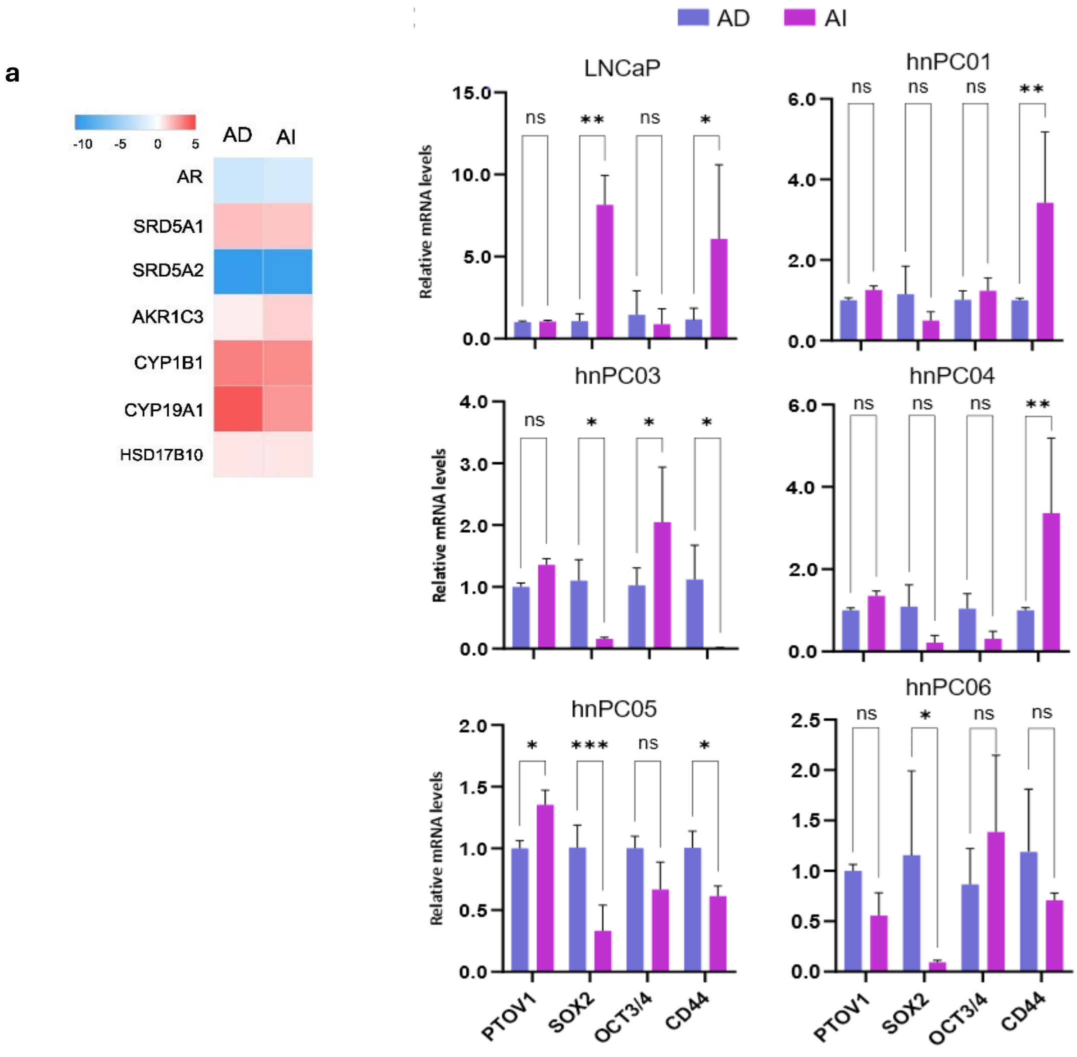

**Figure S6. Analysis of similarity of PCa cell lines, hnPCs and tissues from RP biopsies. a)** Similarity matrix created using Pearson correlation for cell lines (n=4), hnPCs (n=11), and RP tissues (n=11). The correlation coefficients (R) range between -1 (indicating a strong negative correlation) and 1 (indicating a strong positive correlation). **b)** Principal Component Analysis based on gene expression conducted on 26 samples. Formation of two primary clusters is seen: Blue (RP tissues) and yellow (cell cultures). The yellow groups three subclusters. **c)** Heatmap of the expression profiles of hnPCs and original biopsies**.** Relative expressions are shown in Log2 of fold change (BPH) and analyzed in triplicates.

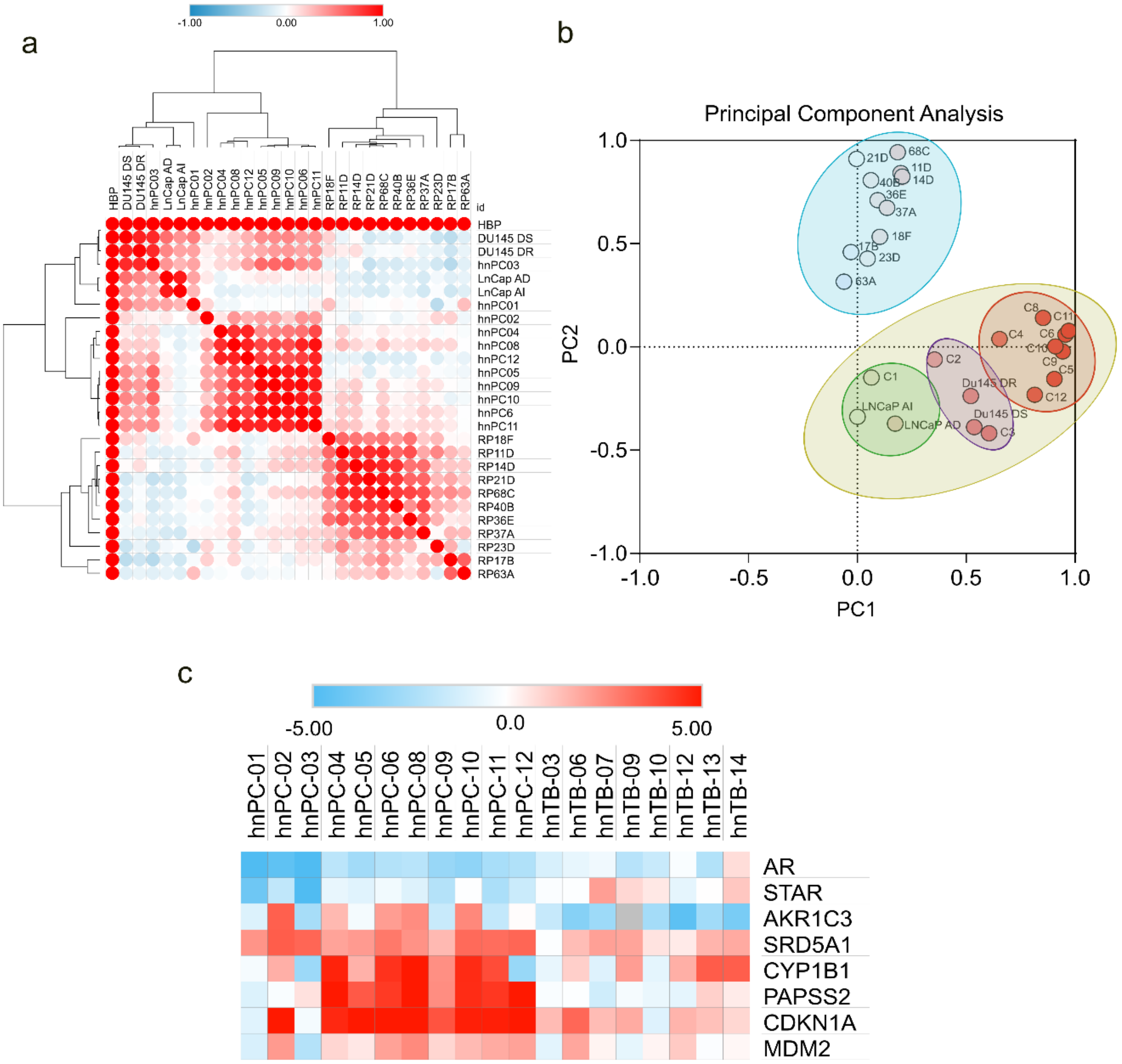

**Figure S7.** Notch1 expression in hnPCs. Notch1 was detected by western blotting in knockdown hnPCs. Experiments were performed in triplicates. Data of protein quantification is the median (+IQR) of the bands area from each siRNA. Protein levels were normalized to β-actin and compared to control.

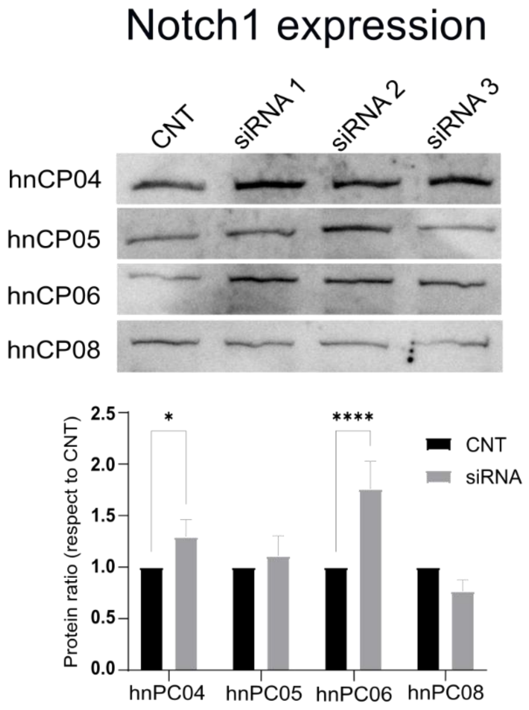
